# Influences of male age, mating history and starvation on female post mating aggression and feeding in *Drosophila*

**DOI:** 10.1101/2020.09.24.311159

**Authors:** Eleanor Bath, Daisy Buzzoni, Toby Ralph, Stuart Wigby, Irem Sepil

## Abstract

1. Mating changes female behaviour and physiology across a wide range of taxa, with important effects for male and female fitness. These changes are often induced by components of the male ejaculate, such as sperm and seminal fluid proteins.
2. However, males can vary significantly in their ejaculates, due to factors such as age, mating history, or nutritional status. This male variation may therefore lead to variation in the strength of responses males can stimulate in females, with alterations in fitness outcomes for both sexes.
3. Using the fruit fly, *Drosophila melanogaster*, we tested whether three aspects of male condition shape an important, but understudied, post-mating response – increased female-female aggression.
4. We found that females mated to old males fought less than females mated to young males. This effect was exacerbated in mates of old, sexually active males, but there was no effect of male starvation status on mating-induced female aggression. There was also a significant effect of age and mating history on female post-mating feeding duration.
5. Our results add to a growing body of literature that variation in male condition can shape sexual selection through post-mating responses in females, including female-female interactions. Studying such variation may therefore be useful for understanding how the condition of sex affects the behaviour of the other.

## Introduction

Mating dramatically alters female behaviour and physiology in a wide range of taxa (McGraw, Suarez, & Wolfner, 2015). It can modify female reproductive rate and output, immune function, activity levels, feeding patterns, and aggression towards conspecifics (Avila, Sirot, LaFlamme, Rubinstein, & Wolfner, 2011; Perry, Sirot, & Wigby, 2013). Mating and associated behavioural changes can also significantly influence female fitness – reducing female lifespan, increasing short-term (but not necessarily long-term) reproductive success, and even affecting offspring survival (Avila et al., 2011; Perry et al., 2013). A number of these mating-induced changes in females are stimulated by components of the male ejaculate. In particular, proteins found in the seminal fluid (seminal fluid proteins – Sfps) have been shown to influence female behaviour and physiology in taxa spanning insects, birds, and mammals, including humans (Avila et al., 2011; Hopkins, Sepil, & Wigby, 2017; McGraw et al., 2015; Ratto et al., 2012). Male ejaculates can therefore significantly affect a female’s immediate and lifetime reproductive output for a wide range of animals, thereby affecting both male and female fitness.

Male ejaculates, however, are not uniform across individuals or time (Bretman, Fricke, & Chapman, 2009; Johnson & Gemmell, 2012; Macartney, Crean, Nakagawa, & Bonduriansky, 2019). Ejaculate component expression appears to be costly, suggesting it is condition-dependent, and males in a population can vary substantially in characteristics that may alter their ejaculate (Linklater, Wertheim, Wigby, & Chapman, 2007; Macartney et al., 2019; Perry et al., 2013). External social and ecological factors can determine how much males invest in their ejaculates. Males in a number of insect species, for example, transfer larger ejaculates or more of specific Sfps to females when rival males are present (Hopkins, Sepil, Bonham, et al., 2019; Kelly & Jennions, 2011). Males in an even broader array of taxa have also been shown to plastically adjust their ejaculate in response to female condition – transferring larger ejaculates to virgin or larger females where a larger ejaculate increases their fertilization success or success in sperm competition with other males (Bonduriansky, 2001; Kelly & Jennions, 2011).

A male’s individual condition is also critical for determining males’ ability to produce and transfer ejaculates. Intrinsic factors such as age, mating activity, and diet have all been shown to influence male ejaculate quality, quantity and composition, impacting both male and female fitness (Fricke, Bretman, & Chapman, 2008; Macartney et al., 2019; Sepil et al., 2020). In both vertebrate and invertebrate taxa, older males tend to transfer smaller or lower quality ejaculates, leading to a decline in fertility or offspring fitness (Dean et al., 2010; Preston, Saint Jalme, Hingrat, Lacroix, & Sorci, 2015; Ruhmann, Koppik, Wolfner, & Fricke, 2018). Similarly, diet-induced changes in male ejaculates are common across taxa. When nutrients are limited, males have both reduced seminal fluid and sperm quantities, leading to reduced fertility or offspring survival (Macartney et al., 2019). Reductions in ejaculate quality mediated by male condition can therefore have important fitness consequences for females (Dean et al., 2010). Conversely, males transferring low quality ejaculates may stimulate weaker post-mating responses in females, reducing the potential harm to females, hence extending female lifespan and increasing female lifetime reproductive success (Koppik & Fricke, 2017; Koppik, Ruhmann, & Fricke, 2018; Ruhmann et al., 2018). Mating-induced changes in females may therefore be highly variable and dependent on both male and female characteristics that shape ejaculate composition, transfer, and storage.

A potentially key but understudied post-mating response is the increase in female-female aggression (hereafter ‘female aggression’) in the fruit fly *Drosophila melanogaster* (Bath et al., 2017; Nilsen, Chan, Huber, & Kravitz, 2004). Across taxa, female aggression can have substantial fitness consequences, with winners in intrasexual aggressive encounters garnering considerable fitness benefits, in the form of increased access to food, better nesting or oviposition sites, protection of their offspring, or social dominance (Clutton-Brock & Huchard, 2013; Rosvall, 2011). In *D. melanogaster*, mated females spend twice as much time engaging in aggressive interactions as virgin females when competing over food (Bath, Biscocho, Easton-Calabria, & Wigby, 2020; Bath et al., 2017; Nilsen et al., 2004). Like other post-mating responses (PMRs), increased female aggression after mating is stimulated by components of the male ejaculate in this species - females must receive sperm to maximally increase aggression post-mating, while the Sfp ‘sex peptide’ is also partially responsible for the increase in aggression after mating (Bath et al., 2017). However, we currently know little about variation in this response. Females have been shown to vary in their aggressive response to ejaculates, with females raised at high larval density showing a greater increase in aggression than those raised at low larval density (Bath, Morimoto, & Wigby, 2018), but we know nothing about how variation in male condition could affect the aggression PMR, despite evidence that male condition significantly alters other female PMRs (Guo & Reinhardt, 2020; Morimoto & Wigby, 2016; Ruhmann et al., 2018). By evaluating whether variation in male traits influences rates of post-mating female aggression, we can begin to discover how inter-sexual interactions that are influenced by the environment can also shape subsequent intrasexual interactions, such as female aggression. Female aggression is therefore a behaviour that is well suited to test if variation in male ejaculates can alter female post-mating responses in a way that influences downstream fitness.

Here we use the fruit fly *Drosophila melanogaster* to test how variation in three male traits known to influence ejaculate characteristics affect an important female post-mating behaviour – female aggression. We chose male age, mating history, and adult feeding status as our male traits, as all three characters are related to male condition and have been shown to strongly influence some aspect of the ejaculate in *D. melanogaster* and other species (Fricke et al., 2008; Macartney et al., 2019; Sepil et al., 2020). In *D. melanogaster*, male reproductive function declines with age, but this is heavily dependent on male sexual activity and differs for different ejaculate components. Sperm production declines with age, regardless of mating history, and old sexually active males transfer fewer sperm to females, leading to severe reductions in fecundity and reproductive success (Koppik & Fricke, 2017; Koppik et al., 2018; Sepil et al., 2020). Surprisingly, sexually active males show no change in their Sfp production and transfer with age. Instead old sexually inactive males transfer a lower abundance of Sfps to females, but their mates show no change in their fecundity or reproductive success (Sepil et al., 2020). Finally, for some Sfps, protein quality declines with age, partly explaining the age-related reduction in male reproductive function. What we know of *D. melanogaster* in the wild suggests that they experience overlapping generations (Behrman, Watson, O’Brien, Heschel, & Schmidt, 2015), hence it is likely that females would come into contact with males of different ages and mating histories. Females may therefore have experienced such variation in male ejaculates throughout their evolutionary history and might have evolved the sensitivity to detect such differences. We therefore investigated male age and mating history together as this enables us to disentangle the effects of both sperm and Sfp declines on rates of post-mating female aggression.

In *Drosophila melanogaster*, nutrition post-eclosion is essential for males to gain sexual maturity (Droney, 1996). Changing the composition of larval and adult diets in males has been shown to alter production and transfer of sperm, success in sperm competition, female reproductive rates, and ability to prevent females from remating in *D. melanogaster* and other insect species (Engels & Sauer, 2007; Guo & Reinhardt, 2020; McGraw et al., 2007; Morimoto & Wigby, 2016). In addition, males starved during adulthood transferred at least one ejaculate component in lower abundance during mating – cis-vaccenyl acetate (cVA) – with the possibility that starvation also affects multiple other ejaculate components (Lebreton et al., 2014). As *D. melanogaster* predominantly eat and lay eggs on rotting fruit (a temporally and spatially patchy resource) (Behrman et al., 2015; Turelli, Coyne, & Prout, 1984), it is possible that males undergo certain periods of time where they face nutrient limitation during adulthood, affecting their ejaculate. Females may therefore come into contact with a range of males that differ in their feeding history and their ejaculate quality and composition.

Given previous studies have already demonstrated the impact of age, mating history and adult nutrition on the ejaculate, we predicted that:

1. Old, sexually active males will stimulate less female aggression than young males as they transfer fewer sperm to females during mating,
2. Old, sexually inactive males will stimulate less aggression than young males, as they transfer a lower abundance of Sfps,
3. Males starved during adulthood will stimulate less aggression than fed males as they transfer a lower abundance of Sfps/smaller ejaculates.

## Methods

### Fly stocks and culture

We used the Dahomey wild-type stock, which was first collected in Benin, Africa in 1970 (Wigby & Chapman, 2004). Flies have been maintained in large, outbred populations in cages with overlapping generations. All flies were cultured and experiments were conducted at 25°C on a 12:12 light: dark cycle in a non-humidified room. Except where stated, adult flies were kept on standard Lewis medium (Lewis, 1960), with no access to live yeast.

Experimental flies were grown at controlled larval density of ~5 larvae per mL of food (~200 larvae in a 75 mL plastic bottle containing ~ 45 mL of food) (Clancy & Kennington, 2001). Flies were collected within 7hr of eclosion to ensure virginity. Females were then kept in individual vials with standard fly medium, but no live yeast.

### Experimental Design

Methods for generating the age and mating history treatments are described in full in Sepil et al., 2020. Briefly, experimental males were placed in groups of 12, consisting of all males (‘unmated’: U) or of 3 virgin males and 9 virgin females (‘frequently-mated’: F). Males from two age classes were used: 1 week (1w, young) and 5 weeks (5w, old) old. Flies in the U treatment were transferred to new vials once a week and those in the F treatment were transferred twice a week using light CO_2_ anesthesia at each transfer. Dead or escaped females were replaced with similarly aged females at each transfer. At 3 weeks, females from the 5-week group were replaced with virgin 3-5 day old females, to reduce co-ageing effects. The males from F were merged into single-sex groups of 10-12 males four/five days before assaying, in order to provide a consistent period of sexual rest prior to the assay point.

To generate the starvation treatments, males were randomly assigned to the ‘fed’ or ‘starved’ treatment upon collection directly after eclosion. Males from the ‘fed’ treatment were kept in vials with standard fly medium, while males in the ‘starved’ treatments were kept in vials containing only damp cotton wool, which provides no nutritional value. Males in both treatments were kept in groups of 15-20 for three days prior to mating. Females were either mated to the ‘fed’ or ‘starved’ males, or were kept as virgins. This experiment was conducted in two blocks.

### Mating assays

Mating assays took place when the experimental females were 3-5 days old. Females were randomly assigned to a treatment group and an individual male was aspirated into each female vial. Pairs were given five hours to mate. Latency to mating and mating duration were recorded for each pair. If pairs did not mate during this time, they were discarded. However unmated males from the 5w-F, sexually active treatment were tested again the following day to increase sample size as such males have been shown to be less likely to mate than the other treatments (Sepil et al., 2020). Only 6 females in our aggression analyses mated with reused males (n = 3 dyads), and results remained the same regardless of whether these 3 dyads were included in analyses or not. We therefore kept these 3 dyads in our main analyses. Once females had mated, they were removed and placed into new vials containing regular fly food medium and no live yeast. We also set up vials with virgin females as controls, where virgin females were switched to new vials at the same time as mated females. Females remained in these vials overnight until they were used in aggression assays. These vials were subsequently frozen to count eggs laid by each female. The mating assays were performed over three days.

### Aggression assays

Females from the same treatment were used in contests 24 hours after mating. For the two hours directly before being used in a contest, females were kept in a vial with damp cotton wool but no food. This brief period of starvation has been shown to increase how much aggressive behaviour females show (Edwards, Rollmann, Morgan, & Mackay, 2006). Flies were then aspirated from these vials into a contest arena (diameter 2 cm) containing an Eppendorf tube cap filled with regular fly food (diameter 5 mm) and a ~2-μl drop of yeast paste, providing a limited resource to fight over. Females were allowed 5 minutes to acclimatise to the arena and were then filmed for 15 minutes using Toshiba Camileo X400 video cameras.

### Behavioural scoring

Videos were scored manually by three different observers using the video analysis software programs JWatcher Video or BORIS (Blumenstein, Evans, & Daniels, 2006; Friard & Gamba, 2016). Videos were scored blind to treatment. We recorded the number of encounters that included headbutts, the duration of each encounter that included headbutts, as well as the amount of time females spent fencing (a lower intensity aggressive behaviour where flies hit each other with their legs). An encounter began when one female headbutted the other and ended when one fly left the food cap, when the flies were one body length apart, or had stopped physically touching each other for three seconds. We also recorded the amount of time each fly spent on the food cap, which we use as a proxy for feeding duration, as we were unable to detect when flies were actually feeding from the food cap.

### Statistical analyses

All analyses were conducted in R (version 3.6.1) (R Core Team, 2012).

#### Aggression

We analysed three measures of aggression –the total duration of time spent in bouts that included headbutts, the number of bouts that included at least one headbutt, and the total amount of time females spent fencing. For headbutt duration, we square root transformed the data to fit the assumptions of a linear model. Headbutt number was analysed using a Generalised Linear Model (GLM) with a quasipoisson distribution as headbutt number was count data that was overdispersed. For the age and mating history experiment, fencing duration fit the assumptions of a linear model with no transformations required. In the starvation experiment, we only collected data on fencing in the second block and fencing duration was analysed using a GLM with a Gamma distribution and inverse link.

In all analyses, the age and mating history experiment data was analysed using two separate models. The first only included mated females so that we could investigate whether it was age, mating history, or an interaction between the two that alter the behaviour. As behavioural assays took place over three days, we also included day as a fixed effect in the model. This model allowed us to investigate which aspects of age and mating history explained differences in post-mating female aggression. The second model had all treatments (including virgins), and included treatment and day as the main explanatory variables. This allowed us to test for differences between females in each of the mating treatments and virgins. For the starvation experiment, we used only one model, with treatment (i.e. ‘fed’, ‘starved’, or virgin), and block included as the main explanatory variables. Results for day (in the age and mating history experiment) and block (in the starvation experiment) are reported in the Supplementary Information.

#### Food cap duration

As most female aggression takes place on the food cap over access to food, we measured the amount of time females spent on the food cap, and use this as a proxy for female feeding duration. In both age and mating history models, a linear model with no transformations was the best fit for the data. In the starvation experiment, the model that best fit the data was a GLM with a Gamma distribution and inverse link. To avoid non-independence of data points, we used dyad as the unit of replication for analyses of contest behaviour and feeding duration, rather than individual flies.

Again in all analyses, to establish the significance of individual factors, we conducted Type II ANOVA tests, using the ‘Anova’ function in the ‘car’ package (Fox & Weisberg, 2019). All multiple comparisons were conducted using a Tukey test from the R package ‘emmeans’ (Lenth, 2018).

#### Mating latency and duration

We analysed latency to mate using Cox proportional hazards models. We analysed the data using the R package ‘survival’ (Therneau, 2015). To analyse mating duration, we used a linear model with mating duration as the response variable.

#### Egg laying

The number of eggs laid by females were analyzed using two GLMs. The first one modeled the presence/absence of nonzero values using a binomial error distribution. The second one modeled the nonzero count data using a Poisson error distribution corrected for overdispersion.

## Results

### Aggression

#### Headbutts

There was a significant effect of male age on headbutts. Females mated to old flies headbutted for less time and fewer times overall than females mated to young flies (Duration LM: F_1,101_ = 5.52, p = 0.02; Number GLM: X^2^_1,103_ = 9.20, p = 0.002; Figure 1.a and S.1.a). There was no effect of mating history (Duration LM: F_1,101_ = 0.67, p = 0.42; Number GLM: X^2^_1,103_ = 0.76, p = 0.38), or an interaction between age and mating history (Duration LM: F_1,101_ = 2.27, p = 0.14; Number GLM: X^2^_1,103_ = 3.39, p = 0.07). When including virgin females, there was a significant effect of treatment when comparing all treatments for both headbutt duration and number (Duration LM: F_4,124_ = 5.65, p < 0.001; Number GLM: X^2^_4,128_ = 31.72, p < 0.001; Figure 1.a and S1.a). This was primarily driven by virgins and females mated to Old-F males headbutting for less time and fewer times than all other treatments (pairwise comparisons in Supplementary Tables 1 and 2).

**Figure 1:**
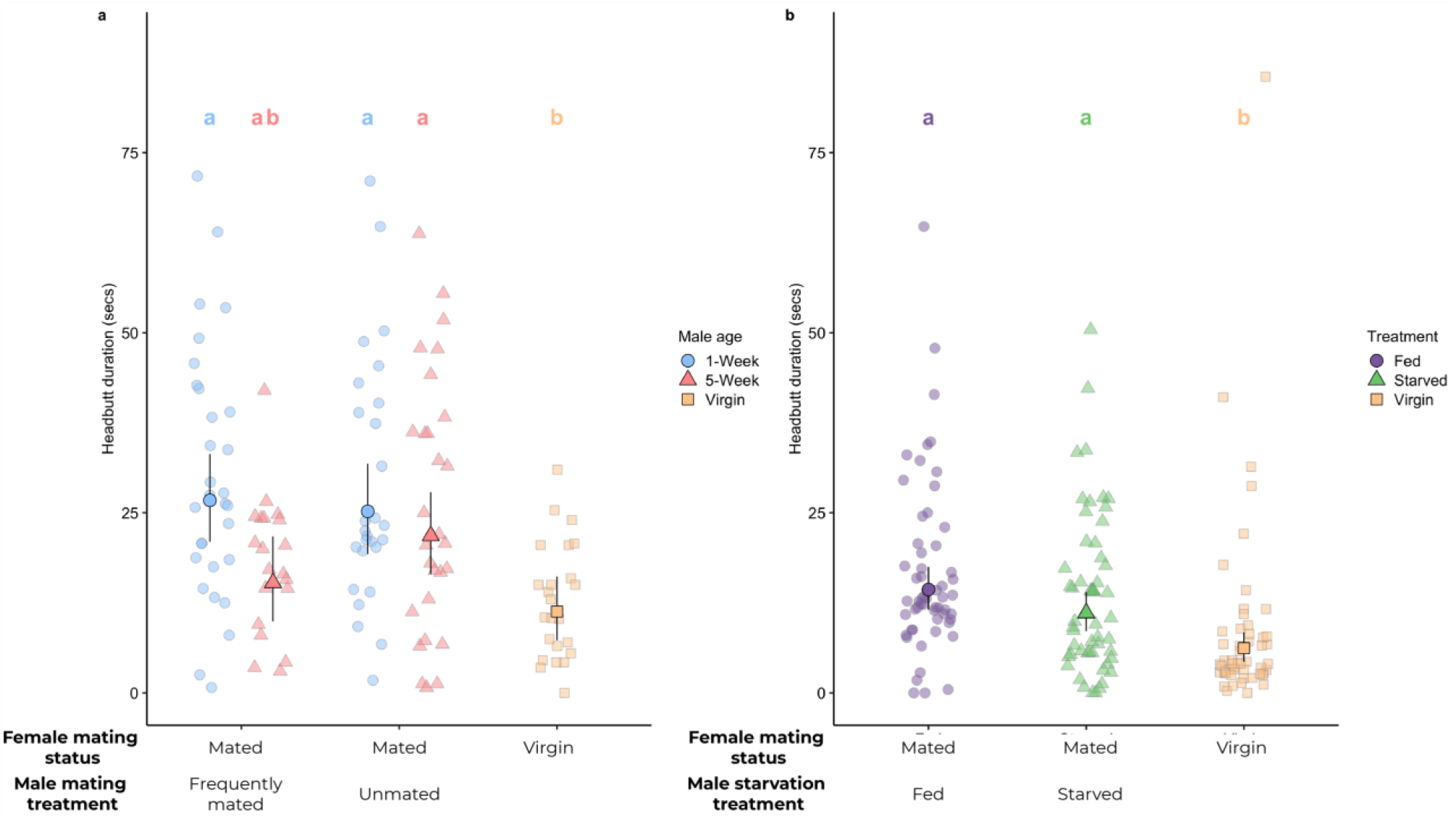
Females mated to old males spent less time headbutting than females mated to young males, but there was no effect of male starvation on female headbutting. **a. Headbutt duration (seconds) by age and mating history** **b. Headbutt duration (seconds) by male starvation treatment** Large points indicate model means (back-transformed to the response scale). Error bars indicate upper and lower 95% confidence limits. Treatments with the same letters were not significantly different from each other using Tukey post-hoc tests.

In the starvation experiment, there was no significant difference between females mated to fed or starved males in headbutt duration or number, but mated females headbutted for longer and displayed more headbutts than virgin females (LM: Duration: F_2,161_ = 10.65, p < 0.001; Number: F_2,161_ = 19.61, p < 0.001; Fig. 1.b and S1.b).

#### Fencing

There was a significant effect of age on fencing duration, with females mated to old males spending less time fencing than those mated to young males (F_1,101_ = 4.72, p = 0.03; Fig. S2.a). There was no effect of mating history (F_1,101_ = 0.14, p = 0.71), or the interaction (F_1,101_ = 0.25, p = 0.61). There was no effect of treatment on fencing duration when looking at all females (F_4,124_ = 1.54, p = 0.2). There was no effect of treatment on fencing duration in the starvation experiment (GLM: X^2^_2,77_ = 2.41, p = 0.3; Fig. S2.b).

### Food cap duration

When looking solely at mated females in the age experiment, there was a significant interaction between age and mating history (F_1,101_ = 6.85, p = 0.01, Fig. 2.a).. When looking at all females, there was a significant effect of treatment (F_4,124_ = 8.41, p < 0.001; Fig. 2.a). Females mated to Old-F males were similar to virgin females, with both spending less time on the food cap than the rest (pairwise comparisons in Supplementary Table 3).

**Figure 2:**
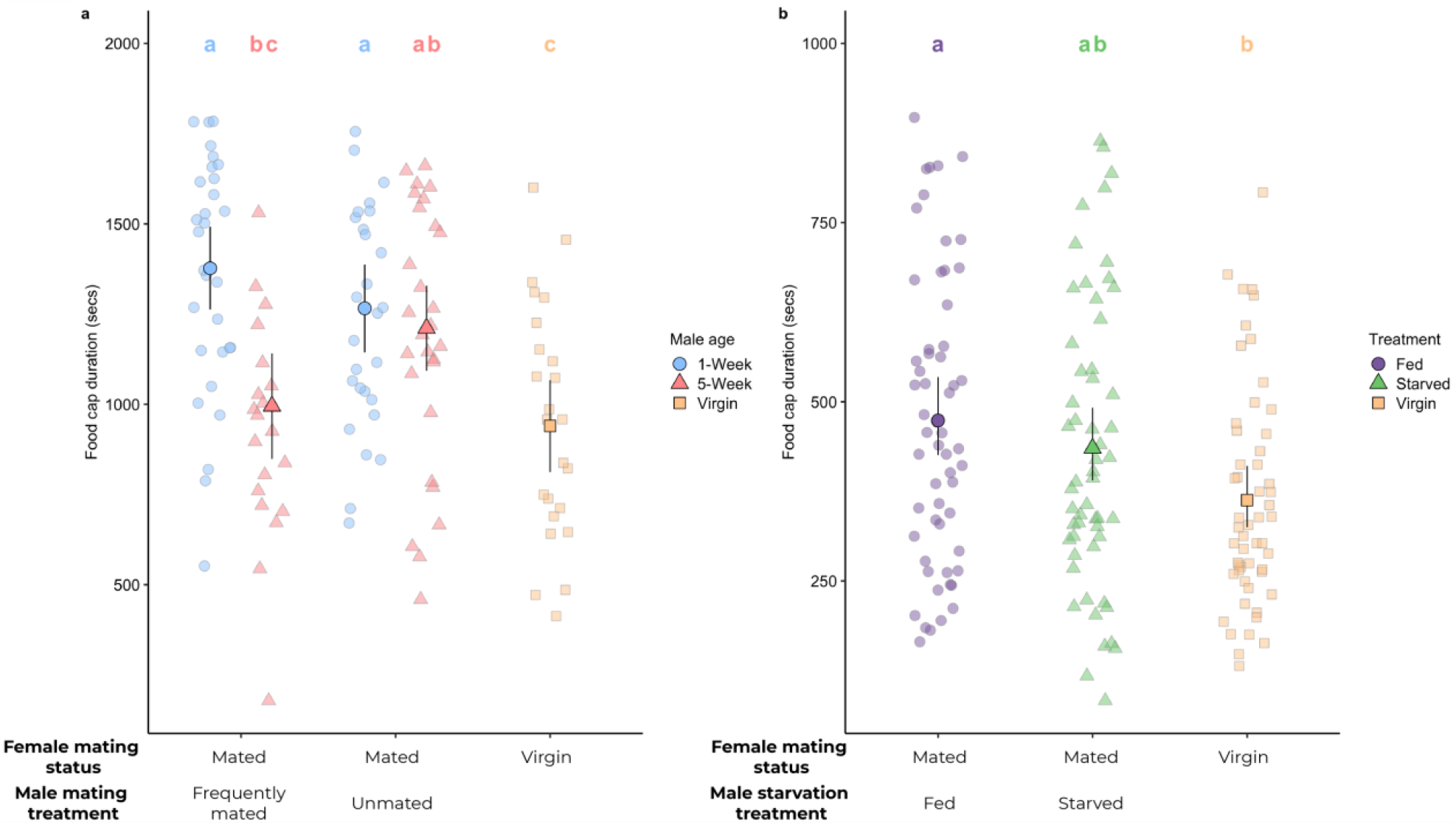
Virgin females and females mated to old males spent less time on the food cap. **a. Food cap duration (seconds) by age and mating history** **b. Food cap duration (seconds) by male starvation treatment** Large points indicate model means (back-transformed to the response scale in b). Error bars indicate upper and lower 95% confidence limits. Treatments with the same letters were not significantly different from each other using Tukey post-hoc tests.

In the starvation experiment, there was a significant effect of treatment on the amount of time females spent on the food cap (X^2^_2,160_ = 10.33, p = 0.006; Fig. 2.b). Females mated to fed males spent longer on the food cap than virgin females (Tukey post-hoc: diff = −0.0006, z = −3.13, p = 0.005), but females in the starved treatment were similar to both virgins and females in the fed treatment (S – V: diff = −0.0004, z = −2.11, p = 0.089; F-S: diff = −0.0002, z = −1.05, p = 0.54).

### Mating latency

Old males and F males were significantly slower and less likely to mate than young and U males (Cox proportional hazards model: Age: X^2^_1,249_ = 16.41, p < 0.001; Mating history: X^2^_1,249_ = 5.2, p = 0.02). This was primarily driven by Old-F males being significantly slower and less likely to mate than all other types of males, as demonstrated by a marginally non-significant interaction between age and mating history (X^2^_1,249_ = 3.71, p = 0.05; Fig. S3.a).

There was no significant difference between fed and starved males in their latency to mate (Cox proportional hazards model: X^2^_1,249_ = 0.04, p = 0.84; Fig. S3.b).

### Mating duration

U males mated for around 4 minutes longer than F males (LM: F_1,254_ = 29.45, p < 0.001; U mean ± SE = 22.9 ± 0.46 mins, F = 19.2 ± 0.49 mins). There was no significant effect of age (F_1,254_ = 0.12, p = 0.73), or an interaction between age and mating history (F_1,254_ = 1.57, p = 0.21).

Females mated to fed males mated for around 3 minutes longer than females mated to starved males (LM: F_1,246_ = 37.88, p < 0.001; Fed mean ± SE = 20.6 ± 0.32 mins, Starved = 17.9 ± 0.32 mins).

### Egg laying

#### Likelihood of laying any eggs

In the age and mating history experiment, we first restricted our analysis to mated females. There were significant effects of both age and mating history on the likelihood of mated females laying any eggs (Age: X^2^_1,261_ = 12.08, p < 0.001; Social: X^2^_1,261_ = 7.82, p = 0.005; Fig. 3.a). Females mated to old males were less likely to lay any eggs, as were females mated to F males. There was no significant interaction between age and mating history (X^2^_1,261_ = 1.41, p = 0.24). When considering all females in the age experiment, there was a significant effect of treatment (X^2^_4,324_ = 96.91, p < 0.001; Fig. 3.a; pairwise comparisons in Table S.4). Virgin females and females mated to Old-F males were less likely to lay any eggs. Only two virgin females laid eggs, while the majority of mated females laid at least one egg, making mated females much more likely to lay eggs overall.

**Figure 3:**
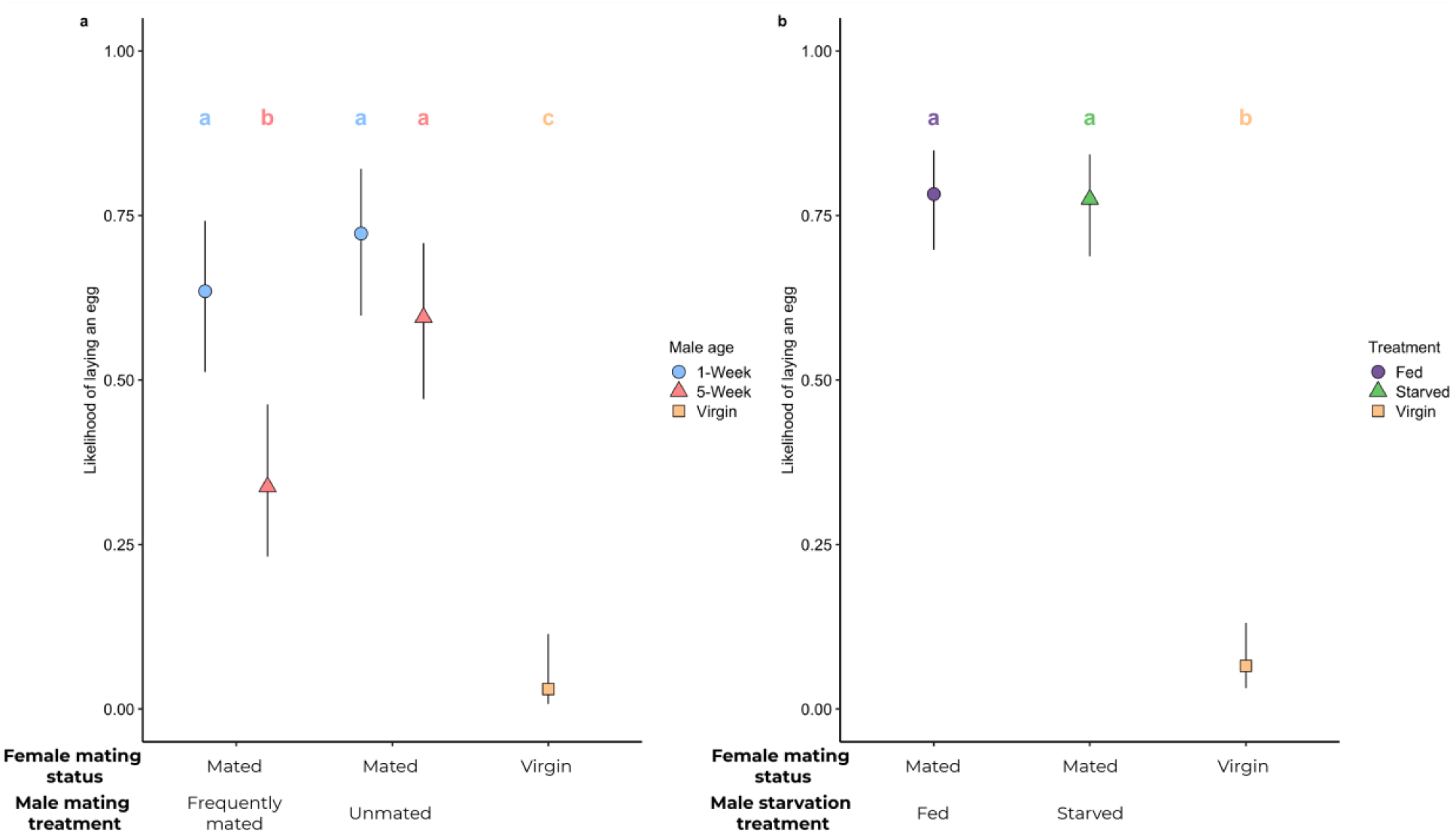
Mated females were more likely to lay eggs than virgins, and females mated to old or F males laid fewer than those mated to young or U males. **a. Likelihood of laying an egg by age and mating history** **b. Likelihood of laying an egg by male starvation treatment** Large points indicate model means (back-transformed to the response scale). Error bars indicate upper and lower 95% confidence limits. Treatments with the same letters were not significantly different from each other using Tukey post-hoc tests.

In the starvation experiment, mated females were more likely to lay eggs than virgins (X^2^_2,327_ = 177.85, p < 0.001, Fig. 3.b), however there was no effect of male feeding status on a female’s likelihood to lay eggs (X^2^_1,1222_ = 0.01, p = 0.93).

#### Number of eggs laid

In the age experiment, when restricting the analysis to just mated females, females mated to old males laid marginally non-significantly fewer eggs (GLM: X^2^_1,261_ = 3.67, p = 0.06). There were no effects of mating history or their interaction (Mating history: X^2^_1,261_ = 0.11, p = 0.74; Interaction: X^2^_1,261_ = 0.01, p = 0.97). When considering all females, there was a significant effect of treatment, where virgin females laid fewer eggs than mated females (GLM: X^2^_4,325_ = 11.02, p = 0.03; Fig. S4.a; pairwise comparisons in Table S.5).

In the starvation experiment, mated females laid more eggs than virgin females in the 24 hours following mating and preceding their aggressive trial (GLM: X^2^_2,327_ = 7.01, p = 0.03; Fig. S4.b). However, there was no significant difference between females mated to fed and starved males (Post-hoc Tukey test: F - S = 0.0008 ± 0.004; z = 0.2, p = 0.98).

## Discussion

We investigated how three aspects of male condition influenced the stimulation of post-mating female aggression. We found that old males stimulated less aggression in females, less time on the food cap, and were less likely to stimulate egg production. This effect of age was particularly strong for mates of old sexually-active males, with females mated to Old-F males being the most similar to virgin females in all our behavioural and fecundity measures.

Our results suggest that these differences in female behaviour due to male condition are likely predominantly driven by changes in Sfp quality, but that age and mating related changes in sperm numbers transferred to females also contribute.

### Male age and mating history

Male age was the main aspect of male condition that affected how much female aggression was stimulated by mating. The quality of some Sfps decline with age, and quantity of sperm declines both with age and mating (Ruhmann et al., 2018; Sepil et al., 2020). These reductions come with major costs – in previous studies, old sexually-active males fathered fewer offspring, were more likely to be infertile, were worse at suppressing female remating, and performed poorly in sperm competition (Koppik & Fricke, 2017; Koppik et al., 2018; Ruhmann et al., 2018; Sepil et al., 2020; Snoke & Promislow, 2003). Our results agree with previous work which indicates females can detect differences among male ejaculates and these differences influence the strength of their PMRs, including aggression (Bath et al., 2017; Bretman, Fricke, Hetherington, Stone, & Chapman, 2010; Fricke et al., 2008; Guo & Reinhardt, 2020; Hopkins, Sepil, Bonham, et al., 2019; Ruhmann et al., 2018; Sepil et al., 2020). Interestingly, different PMRs respond differently to aspects of male variation. While Old-F and Old-U males are both poor sperm competitors, Old-U males do not differ from young males in their reproductive output, fertility, and ability to suppress female remating (Sepil et al., 2020). These differences are primarily due to differences in the ageing of sperm and seminal fluid proteins (Sfps). Sperm transfer declines in Old-F males, but not in Old-U males, but at least some Sfps undergo qualitative changes with age (Sepil et al., 2020). For post-mating female aggression, male age was the only significant factor, with both Old-U and Old-F males stimulating less aggression than young males, suggesting this age difference was potentially driven by a decline in Sfp quality (Sepil et al., 2020). However, females mated to Old-F males showed values closest to virgin females, suggesting a potential role for sperm quantity in determining their level of aggression as well.

Sperm is necessary for females to display a full increase in aggression after mating (Bath et al., 2017). It therefore seems likely that the significant reduction in the amount of sperm transferred to females by Old-F males is primarily responsible for the corresponding reduction in female aggression. Old-F males have previously been shown to transfer both fewer and lower quality sperm (Ruhmann et al., 2018; Sepil et al., 2020), but it is unclear whether it is sperm number, quality, or both, which influence the degree to which mating induces female aggression. Future studies investigating whether it is overall sperm number, number of viable sperm, or Sfp quality that stimulate this aggression would shed light on this matter, as well as the mechanism by which ejaculates stimulate aggression.

We show for the first time that post-mating female feeding duration can also be influenced by male age and mating history. Mates of Old-F males spent the same amount of time on the food cap as virgins, while mates of Old-U males showed the full post-mating increase in feeding duration. These feeding results are consistent with findings that increased female feeding after mating is stimulated by sperm and possibly by Sfps (Carvalho, Kapahi, Anderson, & Benzer, 2006). Feeding and aggression are often linked, with females displaying more aggression also spending more time on the food cap (Bath et al., 2017, 2018). Our results from both experiments suggest that headbutt duration and food cap duration show similar patterns, with females that had been stimulated to spend more time fighting also spending more time on the food cap.

### Male feeding status

How males alter their ejaculate in response to nutrient availability varies dramatically across species. Across both vertebrates and invertebrates, there is a common trend for seminal fluid and sperm quantity to decrease in response to nutrient limitation (Macartney et al., 2019). However, at the same time, there is also a trend in other species for males to increase their investment in ejaculate traits in response to limited nutrients (Mehlis, Rick, & Bakker, 2015; Perry & Rowe, 2010). Adult male *D. melanogaster* with less dietary yeast (a main source of protein) exhibit a reduced ability to prevent females from remating with rival males (a sperm and Sfp-mediated trait) (Fricke et al., 2008). As female aggression is stimulated by both sperm and Sfp transfer, we predicted that it would be a trait sensitive to differences between males in their adult feeding status. However, we found that although male adult feeding status influenced mating duration, it did not significantly alter female aggression or other PMRs, such as egg production. There was a suggestion that females mated to starved males spent slightly less time on the food cap than those mated to fed males, but this was not significant.

Starving males for three days may not have been a strong enough treatment to alter males’ ability to produce or transfer ejaculates in their first mating. However, a similar treatment where males were starved for three days before mating showed a significant reduction in the transfer of a PMR-inducing pheromone (cis-vaccenyl acetate) (Lebreton et al., 2014). We found that starved males mated for around three minutes less than fed males, which suggests a significant effect of starvation on a males’ ability to perform mating behaviours. Mating duration has previously been shown not to correlate with sperm transfer in this species (Gilchrist & Partridge, 2000; Lüpold, Manier, Ala-Honkola, Belote, & Pitnick, 2011), but the relationship between mating duration and the transfer of seminal fluid proteins appears to be more complicated (Hopkins, Sepil, Bonham, et al., 2019; Hopkins, Sepil, Thézénas, et al., 2019; Sepil et al., 2020; Sirot, Wolfner, & Wigby, 2011; Wigby et al., 2009). The differences in mating duration between fed and starved males may indicate differences in the size or amount of ejaculate males were transferring to females, though we found little evidence in the PMRs of females to support this supposition. Males raised in different developmental environments have been shown to transfer similar-sized ejaculates in their first matings despite differences in their ability to produce and replenish them afterwards (Wigby, Perry, Kim, & Sirot, 2016). Starved males may have transferred a larger proportion of their ejaculate reserves during their first mating but be unable to replenish them for a second mating, which we did not test in this experiment. Male feeding status may therefore only begin to influence female PMRs, including aggression, from the second mating onwards.

Whether increased female aggression after mating is adaptive for males or females has yet to be tested. Increasing aggression after mating may be an adaptive response by females as their nutritional needs shift to accommodate their increased investment in reproduction (Boggs, 1981). Acquiring more food, and particularly the more limited high-protein yeast, may require females to compete more vigorously with other females to access limited food resources. Variation in male ejaculate quality or quantity may indicate how much females should increase their overall investment in reproduction, and subsequently, in aggression – i.e. a female mated to an old, sexually active male will receive less sperm and produce fewer offspring and therefore invest less in reproduction than a female mated to a young male. This comparative reduction in reproduction means the first female will require less food and therefore need to show less aggression. Conversely, variation in female aggression due to male ejaculates may not be adaptive but merely a by-product of the means of stimulation of aggression. If aggression is stimulated by the sperm storage organs filling with sperm, for example, then if the sperm storage organs are not filled due to males transferring less sperm, then females will not get as aggressive, even if it would benefit them to do so. Our results are more consistent with the second hypothesis as females mated to Old-F males were less likely to lay any eggs in our experiment, and showed less aggression. However, when they did lay eggs, they laid as many as all other mating treatments, suggesting it is the initial receipt of sperm that is related to the stimulation of aggression.

Potentially, females use mating as a cue to upregulate their aggression. This response may represent more of an ‘on-off’ switch with the receipt of male ejaculate components turning on the aggressive pathway, rather than females increasing their aggression in proportion to the amount of ejaculate they receive. It is possible that females mated to Old-F males did not receive enough ejaculate to switch on this pathway. The idea of an ‘on-off’ switch is consistent with another study which found that different genotypes of males do not stimulate different levels of post-mating aggression in females, despite potential differences in their ejaculates (Bath et al., 2020).

## Conclusion

Variation amongst males can be caused by a number of intrinsic and extrinsic factors. The effects this variation has on males and their mates can shape sexual selection and sexual conflict through mating and the post-mating responses stimulated in females. Here we show that male age can modify the aggression females direct towards other females, potentially altering their access to food and overall fitness. Females across species show plasticity in their aggressive and other reproductive behaviours, often responding to male cues. Understanding how variation in males alters these cues and how females respond to them will improve our understanding of female behaviour and its subsequent fitness effects. Female aggression is closely linked to reproduction and mating across a broad range of taxa, with aggression towards conspecifics increasing dramatically after mating, or in association with key reproductive events (Bath et al., 2017; Palanza, Re, Mainardi, Brain, & Parmigiani, 1996; Seebacher, Ward, & Wilson, 2013). Not only are there clear temporal associations between reproduction and aggression across species, but there is also evidence that some of these changes in female aggression are stimulated by the receipt or perception of various male signals, including auditory, chemical or hormonal signals (Bowler, Cushing, & Carter, 2002; Kapusta & Marchlewska-Koj, 1998; Rillich, Buhl, Schildberger, & Stevenson, 2009). Understanding variation in males and their signals may therefore be critical in understanding variation in female intrasexual behaviours, such as aggression, and their fitness consequences.

## Author contributions

E.B., I.S., S.W. conceived the project. E.B. and I.S. designed the experiments, with contributions from S.W., D.B. & T.R.. E.B., D.B., T.R. & I.S. performed the experiments. E.B., D.B., & T.R. scored the behavioural data and analysed the data. E.B. wrote the manuscript, with editing contributed by D.B., T.R., S.W. & I.S.

## Acknowledgements

This study was funded by a John Fell Fund Grant, University of Oxford (161/126) and a Junior Research Fellowship, Christ Church College, University of Oxford to E.B. S.W. and I.S were funded by a BBSRC fellowship to S.W. (BB/K014544/1). I.S was funded by a BBSRC fellowship (BB/T008881/1).

## Data availability

All data will be made publicly accessible on the Oxford University Research Archive (doi to be inserted here).

## Notes

### Competing Interest Statement

The authors have declared no competing interest.

