## Supplementary Information for "Influences of male age, mating history and starvation on female post mating aggression and feeding in *Drosophila*"

**Supplementary Table 1: Pairwise comparisons for headbutt number in ageing experiment**

Significant values are indicated in **bold, red** text.

| | | Frequently mated | | | | | | Unmated | | | | | |
|---|---|---|---|---|---|---|---|---|---|---|---|---|---|
| | | 1-Week | | | 5-Week | | | 1-Week | | | 5-Week | | |
| **Mating history** | **Age** | Difference | Z | P-value | Difference | Z | P-value | Difference | Z | P-value | Difference | Z | P-value |
| **Frequently mated** | **1-Week** | - | - | - | | | | | | | | | |
| | **5-Week** | 0.56 | 3.3 | **0.009** | - | - | - | | | | | | |
| **Unmated** | **1-Week** | 0.05 | 0.41 | 0.994 | 0.5 | 2.98 | **0.024** | - | - | - | | | |
| | **5-Week** | 0.21 | 1.56 | 0.528 | -0.34 | -1.96 | 0.288 | 0.16 | 1.14 | 0.78 | - | - | - |
| **Virgin** | | 0.76 | 4.46 | **< 0.001** | 0.2 | 0.99 | 0.86 | 0.7 | 4.09 | **< 0.001** | 0.54 | 3.08 | **0.018** |

**Supplementary Table 2: Pairwise comparisons for headbutt duration in ageing experiment**

Significant values are indicated in **bold, red** text.

| | | Frequently mated | | | | | | Unmated | | | | | |
|---|---|---|---|---|---|---|---|---|---|---|---|---|---|
| | | 1-Week | | | 5-Week | | | 1-Week | | | 5-Week | | |
| Mating history | Age | Difference | t | P-value | Difference | t | P-value | Difference | t | P-value | Difference | t | P-value |
| Frequently mated | 1-Week | - | - | - | | | | | | | | | |
| | 5-Week | 1.27 | 2.64 | 0.07 | - | - | - | | | | | | |
| Unmated | 1-Week | 0.16 | 0.36 | 0.99 | 1.11 | 2.29 | 0.16 | - | - | - | | | |
| | 5-Week | 0.5 | 1.17 | 0.77 | -0.77 | -1.58 | 0.51 | 0.34 | 0.78 | 0.94 | - | - | - |
| Virgin | | 1.81 | 4.05 | **< 0.001** | 0.55 | 1.07 | 0.82 | 1.66 | 3.59 | **0.004** | 1.32 | 2.9 | **0.04** |

**Supplementary Table 3: Pairwise comparisons for time spent on the food cap in ageing experiment**

Significant values are indicated in **bold, red** text.

| | | Frequently mated | | | | | | Unmated | | | | | |
|---|---|---|---|---|---|---|---|---|---|---|---|---|---|
| | | 1-Week | | | 5-Week | | | 1-Week | | | 5-Week | | |
| Mating history | Age | Difference | t | P-value | Difference | t | P-value | Difference | t | P-value | Difference | t | P-value |
| Frequently mated | 1-Week | - | - | - | | | | | | | | | |
| | 5-Week | 382.3 | 4.13 | **< 0.001** | - | - | - | | | | | | |
| Unmated | 1-Week | 111.9 | 1.34 | 0.67 | 270.4 | 2.89 | **0.037** | - | - | - | | | |
| | 5-Week | 166.6 | 2.02 | 0.26 | -215.6 | -2.3 | 0.15 | 54.8 | 0.65 | 0.97 | - | - | - |
| Virgin | | 437.8 | 5.07 | **< 0.001** | 55.6 | 0.56 | 0.98 | 326 | 3.66 | **0.003** | 271.2 | 3.09 | **0.02** |

**Supplementary Table 4: Pairwise comparisons for likelihood of laying eggs**

Significant values are indicated in **bold, red** text.

| | | Frequently mated | | | | | | Unmated | | | | | |
|---|---|---|---|---|---|---|---|---|---|---|---|---|---|
| | | 1-Week | | | 5-Week | | | 1-Week | | | 5-Week | | |
| **Mating history** | **Age** | Odds ratio | z | P-value | Odds ratio | t | P-value | Odds ratio | t | P-value | Odds ratio | t | P-value |
| **Frequently mated** | **1-Week** | - | - | - | | | | | | | | | |
| | **5-Week** | 3.38 | 3.32 | **0.008** | - | - | - | | | | | | |
| **Unmated** | **1-Week** | 0.67 | -1.04 | 0.84 | 5.02 | 4.15 | **< 0.001** | - | - | - | | | |
| | **5-Week** | 1.19 | 0.49 | 0.99 | 0.35 | -2.86 | **0.035** | 1.77 | 1.5 | 0.57 | - | - | - |
| **Virgin** | | 55.67 | 5.27 | **< 0.001** | 16.49 | 3.67 | **0.002** | 82.83 | 5.71 | **< 0.001** | 46.73 | 5.04 | **< 0.001** |

**Supplementary Table 5: Pairwise comparisons for number of eggs laid**

Significant values are indicated in **bold, red** text. As only 2 virgin females laid eggs, pairwise comparisons between virgins and other treatment females have a low sample size.

| | | Frequently mated | | | | | | Unmated | | | | | |
|---|---|---|---|---|---|---|---|---|---|---|---|---|---|
| | | 1-Week | | | 5-Week | | | 1-Week | | | 5-Week | | |
| **Mating history** | **Age** | Difference | z | P-value | Difference | t | P-value | Difference | t | P-value | Difference | t | P-value |
| **Frequently mated** | **1-Week** | - | - | - | | | | | | | | | |
| | **5-Week** | 0.98 | -0.02 | 1 | - | - | - | | | | | | |
| **Unmated** | **1-Week** | 0.97 | -0.23 | 0.99 | 1.025 | 0.17 | 0.99 | - | - | - | | | |
| | **5-Week** | 0.8 | -1.82 | 0.36 | 0.81 | -1.49 | 0.57 | 0.83 | -1.62 | 0.48 | - | - | - |
| **Virgin** | | 5.65 | 1.76 | 0.4 | 5.66 | 1.76 | 0.39 | 5.81 | 1.79 | 0.38 | 7.04 | 1.99 | 0.27 |

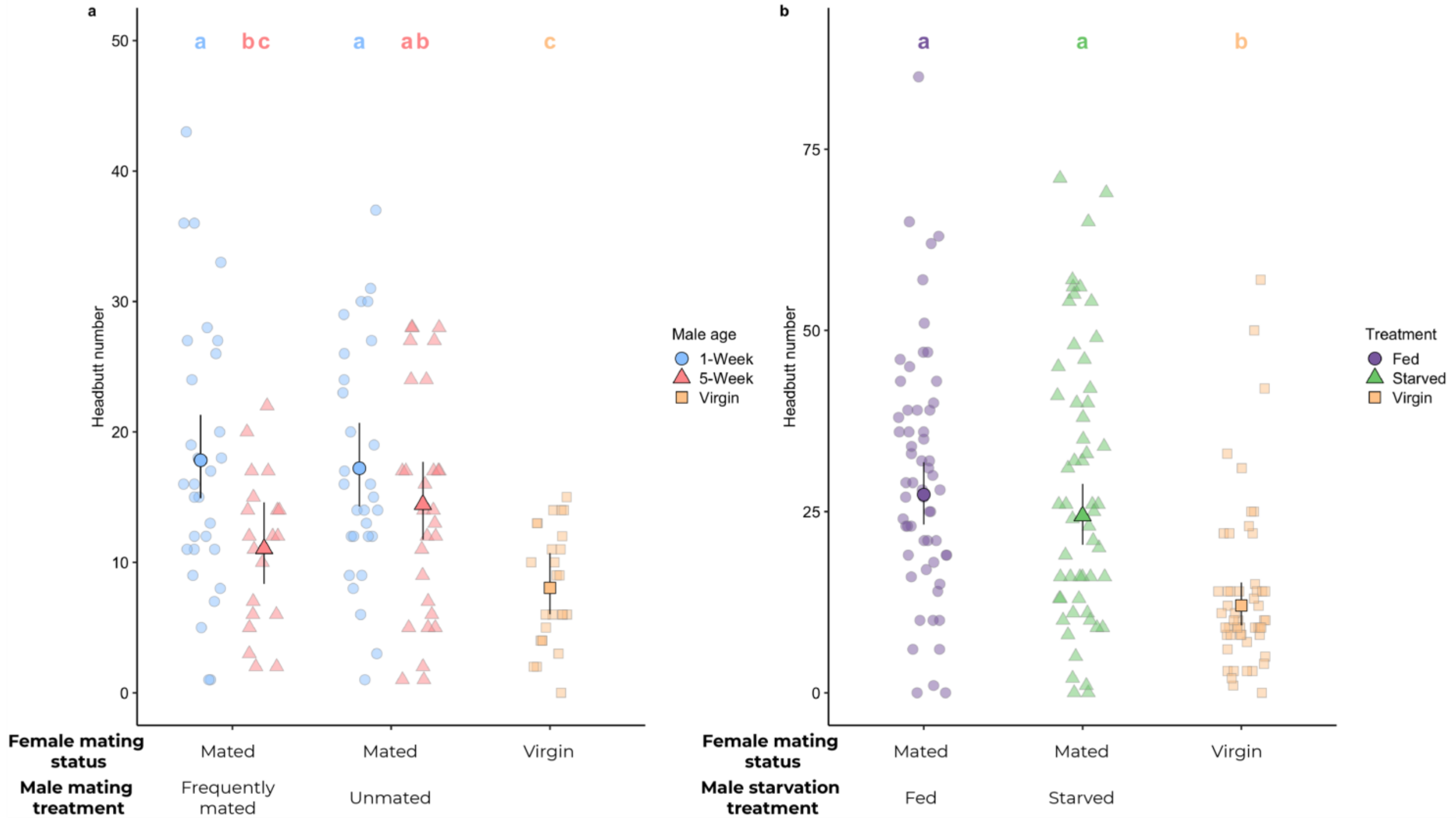


**Supplementary Figure 1: Females mated to old males headbutted less often than females mated to young males**

**a. Headbutt number by age and mating history**

**b. Headbutt number by male starvation treatment**

Large points indicate model means (back-transformed to the response scale). Error bars indicate upper and lower 95% confidence limits.

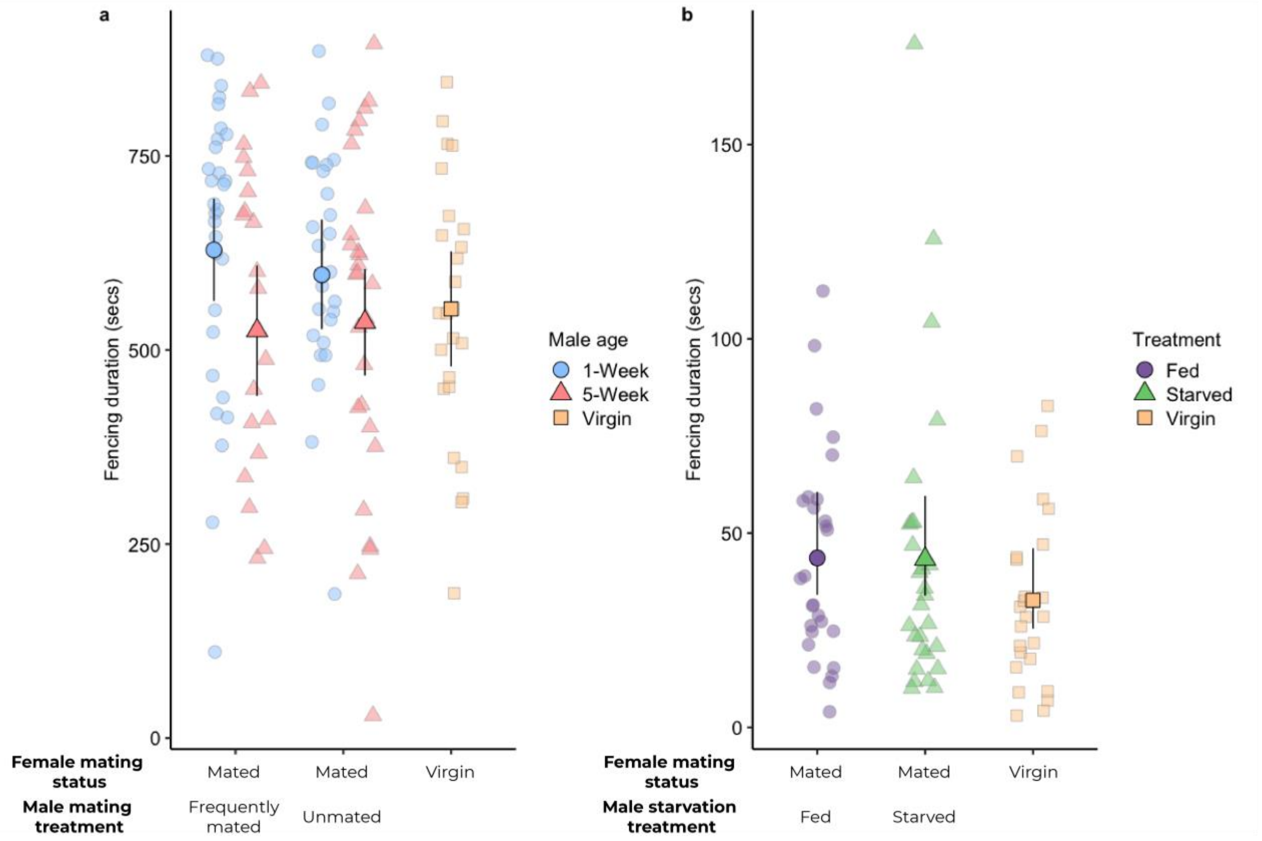


**Supplementary Figure 2: Females mated to old males spent less time fencing than females mated to young males**

**a. Fencing duration (seconds) by age and social history**

**b. Fencing duration (seconds) by male starvation treatment**

Large points indicate model means (back-transformed to the response scale in b). Error bars indicate upper and lower 95% confidence limits.

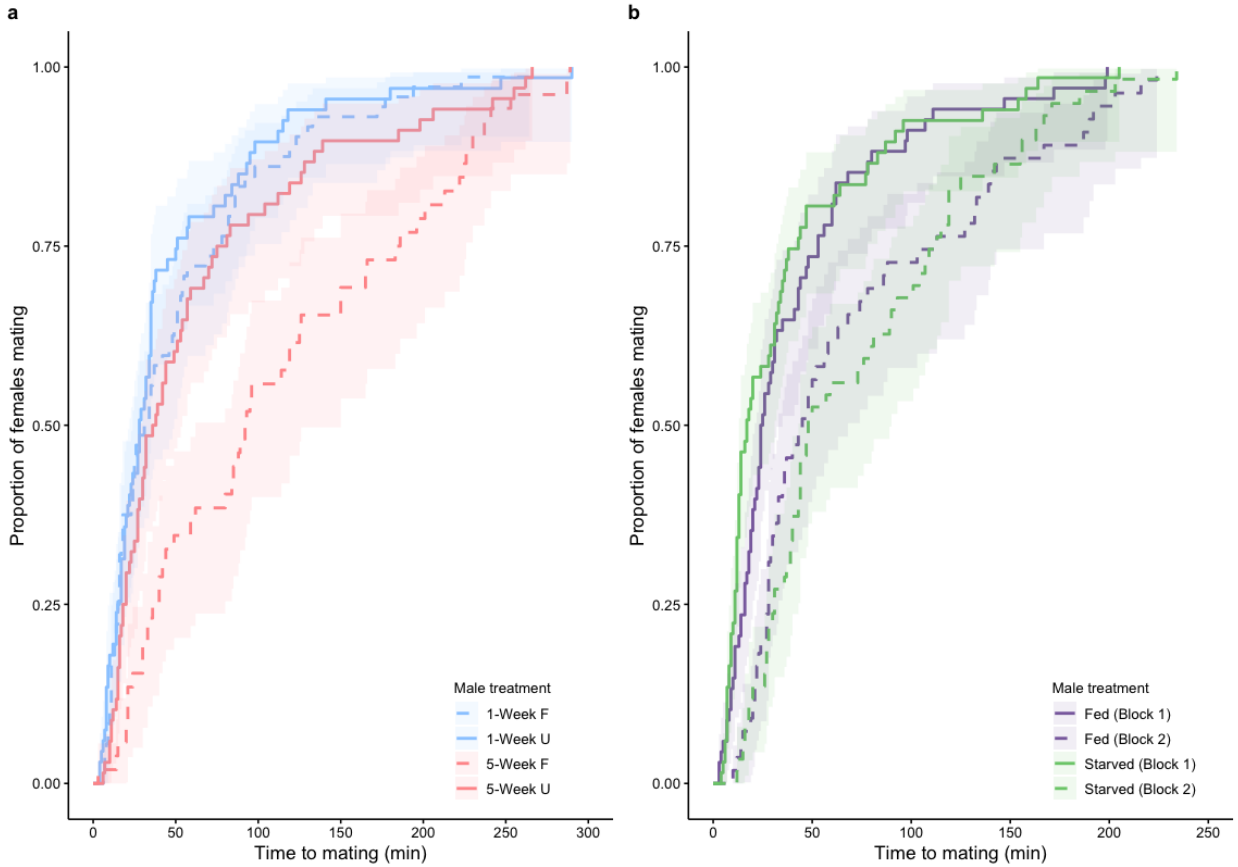


**Supplementary Figure 3: Old and frequently-mated males took longer to mate than young and unmated males, while there was no difference between fed and starved males**

**a. Proportion of females mating by age and mating history**

Red lines indicate old (5-week old) males, while blue lines represent young (1-week old). Dashed lines represent frequently-mated males, while solid lines represent unmated males.

**b. Proportion of females mating by male starvation treatment**

Purple lines indicate fed males, while green lines indicate starved males. Solid lines indicate flies from Block 1, while dashed lines indicate flies from Block 2. The shaded areas in both figures represent 95% confidence intervals.

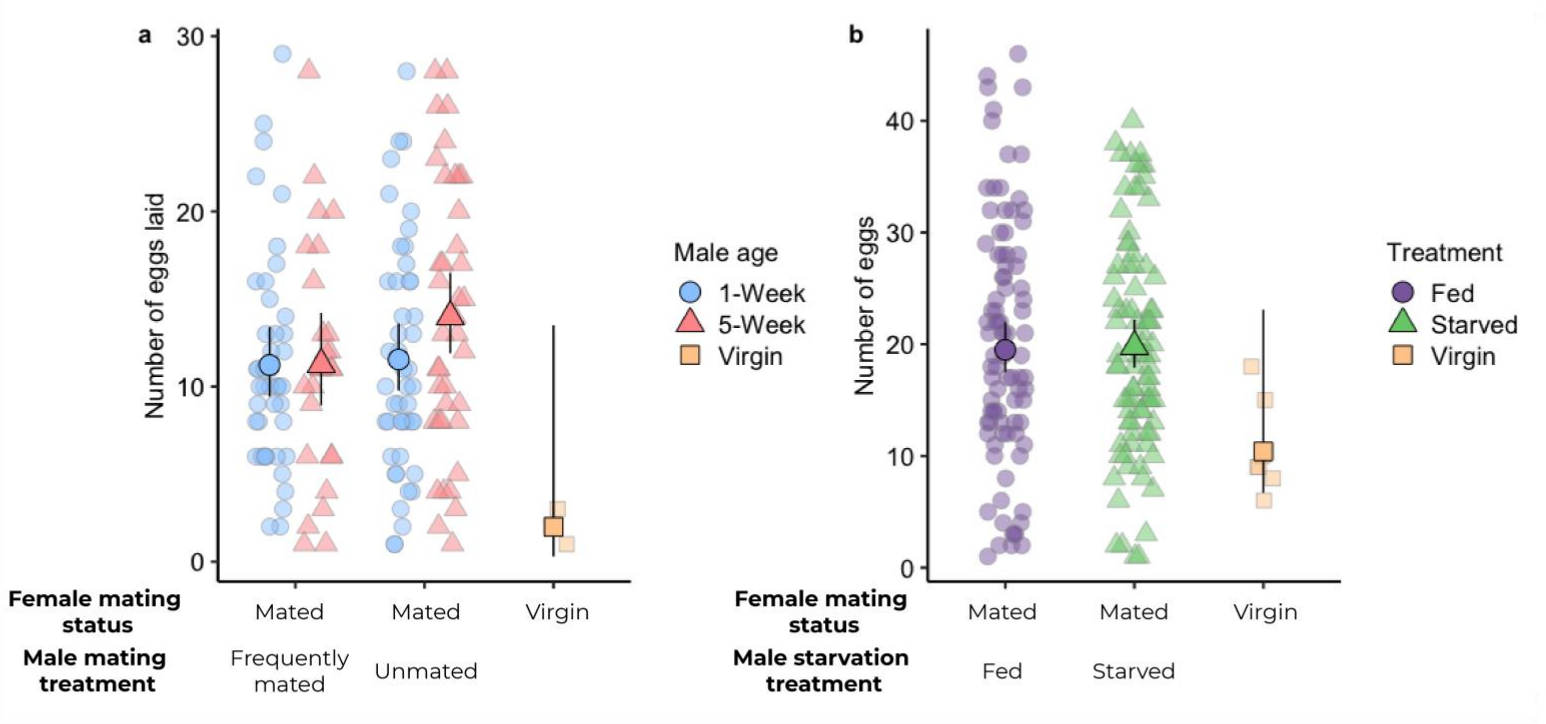


**Supplementary Figure 4: Mated females laid more eggs than virgin females, but there was no effect of male condition**

**a. Number of eggs laid by age and mating history**

**b. Number of eggs laid by male starvation treatment**

Large points indicate model means (back-transformed to the response scale). Error bars indicate upper and lower 95% confidence limits.

### **Block and Day results**

Aggregation

*Headbutts*

In the age experiment, when only mated females were included in the analysis, females headbutted more often on day 1 than subsequent days ($X^2_{2,103}$ = 8.85, p = 0.01), but there was no difference in headbutt duration across days ($F_{2,101}$ = 1.97, p = 0.15). When virgin females were added to the analysis, females again headbutted more often on day 1 than subsequent days ($X^2_{2,128}$ = 6.99, p = 0.03), but there was no difference in headbutt duration across days ($F_{2,128}$ = 0.96, p = 0.39).

In the starvation experiment, females in block 2 headbutted for longer and more often than females in block 1 (Number: $F_{1,161}$ = 38.52, p < 0.001; Duration: $F_{1,161}$ = 43.07, p < 0.001).

*Fencing*

In the age experiment, there was no effect of day on fencing duration for the model with only mated females ($F_{1,101}$ = 1.45, p = 0.24). There was a marginally non-significant effect of day when including all females in the model ($F_{2,124}$ = 2.8, p = 0.06).

We only measured fencing duration in block 2 of the starvation experiment, so cannot test whether there was an effect of block on fencing duration.

Food cap duration

In the age experiment, when only mated females were included in the model, there was a significant main effect of day ($F_{2,101}$ = 3.12, p = 0.049). There was also a significant effect of day when all females were included in the model ($F_{1,123}$ = 8.54, p = 0.004).

In the starvation experiment, there was no effect of block on the amount of time females spent on the food cap ($X^2_{2,160}$ = 0.94, p = 0.33).

Mating latency

In the age experiment, there was no effect of day on mating latency ($X^2_{1,249} = 0.19$, $p = 0.67$).

In the starvation experiment, flies were faster to mate in block 1 than block 2 ($X^2_{1,249} = 27.44$, $p < 0.001$).

Mating duration

In the age experiment, there was no significant effect of day on mating duration ($F_{1,254} = 2.76$, $p = 0.09$).

In the starvation experiment, flies from block 1 showed a tendency to mate for almost a minute longer than flies from block 2 ($F_{1,246} = 4.26$, $p = 0.07$; B1 mean ± SE = 19.6 ± 0.31 mins, B2 = 18.8 ± 0.34 mins).

Egg laying

*Likelihood of laying any eggs*

In the age experiment, when considering only mated females, there was a marginally non-significant effect of day ($X^2_{1,261} = 3.05$, $p = 0.08$). Similarly, when including all females in the model, there was a marginally non-significant effect of day ($X^2_{2,324} = 5.38$, $p = 0.07$).

In the starvation experiment, females from block 1 were more likely to lay eggs than females in block 2 ($X^2_{1,222} = 17.52$, $p < 0.001$).

*Number of eggs laid*

In the age experiment, when considering only mated females, there was no effect of day on the number of eggs laid ($X^2_{1,261} = 0.09$, $p = 0.76$). Similarly, there was no effect of day when considering all females ($X^2_{1,325} = 0.02$, $p = 0.88$).

In the starvation experiment, there was no effect of block on the number of eggs laid ($X^2_{1,237} = 1.54$, $p = 0.21$).
